# Antennal RNAseq reveals odorant receptors with sex-biased expression in the common eastern firefly, *Photinus pyralis*

**DOI:** 10.1101/2025.03.21.644631

**Authors:** Sarah E. Lower, Samuel Pring, Hanh Tran, Katherine Martinson, Susan Deering, Mathew A. Price, Brian Vestal, Gregory M. Pask, Douglas B. Collins, Robert F. Mitchell

**Author notes:** **Corresponding author**: Sarah Lower.

## Abstract

**Background:** With their charismatic nighttime flashes, fireflies are a classic organismal system for studying visual mating signal evolution. However, across their diversity, fireflies employ a variety of mating strategies that include both chemical and visual signals. While phylogenetic evidence points to a common ancestor that relied on long-range pheromones, behavioral evidence suggests that light-dependent flashing fireflies do not use smell for mating. We tested this hypothesis by investigating the olfactory genetics of the nocturnal, light-using common eastern firefly, *Photinus pyralis.* In insects, odors are primarily detected by odorant receptor (OR) proteins embedded in the dendritic membranes of olfactory receptor neurons. If pheromones are part of mate signaling in light-using fireflies, then one or more OR genes should be upregulated in the antennae of the searching sex (males). We therefore annotated the complete suite of ORs from the genome of *P*. *pyralis* and measured expression of OR genes between the sexes.

**Results:** We identified 102 ORs in the *Photinus pyralis* genome, including the conserved single-copy Orco. Our phylogenetic analysis showed lineage-specific OR diversification in *P. pyralis* relative to other beetle species. Differential expression analysis of male and female antennae and hind legs revealed that a subset of ORs are upregulated in antennae as compared to legs, suggesting a role in adult olfaction. Notably, PpyrOR6 was one of two genes, and the only OR, that was significantly upregulated between male and female antennae, suggesting a role in mating.

**Conclusions:** These findings increase known diversity of insect ORs in an understudied beetle family and suggest that bioluminescent fireflies use multimodal signals during mating.

## Background

Fireflies, in the beetle family Lampyridae, are renowned for their nocturnal bioluminescent signals that adults use to recognize, find, and choose mates. However, some species lack light-producing organs as adults, are diurnal, and rely on volatile pheromones for mating [1]. Some species even use both dim glows and pheromones [2–4]. This mating signal diversity makes fireflies ideal for studying chemosensory evolution.

While light signals are clearly important for mating in nocturnal species, less attention has been paid to chemical signals. Phylogenetic studies suggest that the ancestor of all fireflies was diurnal and non-luminescent and instead used pheromones for communication [5–7]. While all firefly larvae glow, supporting a single origin of bioluminescence early in firefly evolution, adult bioluminescence subsequently evolved either one or several times, with multiple secondary losses of light and reversion to pheromones occurring independently across firefly lineages. The evolutionary lability of mating signal mode in fireflies suggests that the molecular machinery for pheromone signaling and reception during mating remains, either functional or not, in light-using lineages. However, behavioral research using hexane-washed freeze-killed females suggests that at least close-range, cuticular chemical signals are important for species recognition only in non-luminescent species [8].

Fireflies, like other insects, likely detect odors primarily using their antennae, which are covered in sensilla that contain olfactory receptor neurons (ORNs; reviewed in [9]). Odor molecules enter through pores in the sensilla, and either diffuse through the sensillar lymph or are transported by proteins to the ORN dendrites where they bind odorant-gated ion channels that open and induce an action potential. The most common receptors mediating insect olfaction are the odorant receptors (ORs), which are heterotetrameric complexes of an odorant receptor co-receptor, Orco, and variable odorant receptor, OrX, the latter of which confers odorant specificity [10,11]. All ORs share a basic structure with seven transmembrane domains and an intracellular amino-terminus; however, OrXs show high copy number and sequence diversity across insect groups, while Orco is single-copy and conserved in sequence across all insect species examined to date [12].

High-throughput sequencing has facilitated the exploration of the molecular basis of chemosensation in diverse non-model insect lineages. Complete OR repertoires have now been annotated from many exemplar species within the largest insect orders, revealing species-specific OR diversity driven by rapid evolution [13–15]. For example, beetles have anywhere from tens to hundreds of functional ORs that are organized into seven major clades [14], with many copies departing from the canonical 5-exon (A,B,C,D,E) structure due to intron insertions and/or deletions. Nevertheless, even well-studied insect orders remain undersampled; for example, ORs from the diverse beetle infraorder Elateriformia, which includes fireflies, are described from only a single species, *Agrilus planipennis* (Buprestidae; [21]), which is a dietary specialist with a reduced OR repertoire.

Here, we investigated OR evolution in *Photinus pyralis*, a well-studied firefly species for its nocturnal light-based mating displays (e.g., [16–18]). Using comprehensive manual annotation, we curated the full OR repertoire of *P. pyralis* and explored its diversification in structure and location throughout the genome. Further, using phylogenetic analysis, we investigated *P. pyralis* OR evolution relative to other beetle ORs. Finally, we examined OR expression using RNA sequencing of adult male and female antennae and hind legs. We found that *P. pyralis* hosts a large OR repertoire, with a dramatic Group 2B expansion found on 10 out of 11 linkage groups, including the putative sex chromosome. A third of ORs were significantly upregulated in male and female antennae as compared to legs, with a single OR, PpyrOR6, upregulated in male antennae as compared to female antennae. Taken together, these results provide a foundation for studying the evolution of olfaction in fireflies, and they suggest that males of light-using firefly species rely on olfaction for mating more than previously thought.

## Methods

### OR identification

We identified putative OR nucleotide sequences in the published *Photinus pyralis* genome assembly (GCA_008802855.1; [18]) using an exhaustive iterative tblastn strategy followed by manual curation [14]. Briefly, for searchability, we divided genomic scaffolds into 200 kb contigs, then we determined which contigs contained OR-like sequences using tblastn (e-value: 1.0; [19]). We used 547 ORs from six beetle species for tblastn search and subsequent annotation, selected to span the phylogeny of beetles and represent all beetle OR subfamilies: *Anoplophora glabripennis*, N = 132 [20]; *Agrilus planipennis*, N = 46 [21]; *Hydroscapha redfordi*, N = 39 [14]; *Nicrophorus vespilloides*, N = 84 [14]; *Priacma serrata*, N = 123 [14]; and *Rhyzopertha dominica*, N = 123 [22]. We imported promising contigs into Geneious Prime v. 2021.2 (https://www.geneious.com) for manual curation. We annotated exons and appropriate splice sites by repeated blastx (e-value: 1.0) of contig sequence against the database of known beetle ORs. We named ORs with missing C-terminal or N-terminal sequence, or including null mutations (pseudogenes), with suffixes CTE, NTE, and PSE, respectively. We then reiterated the process using newly-identified ORs as queries until no new ORs were identified. Finally, we compared ORs to those predicted from the most recent genome assembly (v1.3; GCF_008802855; [18]) and incorporated any additional sequence data from the predictions into our models. Where possible, we confirmed OR sequence and splice junctions using Illumina RNA sequencing reads from the antennae of one male (SEL534) and one female (SEL672) *P. pyralis* (see RNA extraction and sequencing below) mapped onto the contigs containing manually annotated ORs using the Geneious RNA read mapper (parameters: find novel introns up to 30,000 bp).

To further support the annotation, we predicted transmembrane domains for all OR models, including full length, partial, and pseudogenized genes, using TOPCONS, a widely-used membrane topology prediction web server (https://topcons.cbr.su.se/pred/; [23]). The TOPCONS algorithm combines topology predictions from five sub-methods into a single topology profile, and has been shown to be equally or more accurate for membrane topology prediction than other software [23].

To ensure that newly identified OR names were informative as possible, after removing stop codons and ambiguous amino acids, we compared all *P. pyralis* OR (PpyrOR) amino acid sequences, including CTE, NTE, and PSE sequences, with OR amino acid sequences from three other representative beetle species with well-curated OR annotations, excluding pseudogenes and sequences shorter than 100 amino acids: *A. plannipennis*, the sole elateriform relative of fireflies where genomic ORs have been described [14,24]; *N. vespilloides*, the closest polyphagan relative to Elateriformia with annotated ORs [14], and *Tribolium castaneum*, a model beetle, with group 5 ORs otherwise lacking from *A*. *planipennis* and *N*. *vespilloides* [14,25]. We built an initial phylogeny for naming PpyrORs by aligning these sequences with MUSCLE v. 5.3 (parameters: default; [26]), then trimming the resulting alignment to conserved regions using trimAL v. 1.5.0 (parameters: automated1; [27]), constructed a phylogeny using FastTree v. 2.1.11 (parameters: default; [28,29]). We visualized the resulting tree in Geneious Prime. To incorporate evolutionary and genomic location information, where possible we named PpyrORs sequentially according to their position in the phylogeny relative to known beetle ORs, and following gene order within tandem arrays.

### Phylogeny

To more rigorously examine PpyrOR diversification and place it in context of potential OR function, we next constructed a maximum likelihood tree that included testing for a model of evolution and 13 additional beetle ORs with known ligands (Additional File 2: Table S1). Using Geneious Prime, we aligned amino acid sequences with MUSCLE 5.1 (parameters: algorithm PPP; [30]). We subsequently used Noisy v. 1.5.12.1 [31] on NGPhlyogeny.fr (default parameters; [32]) to identify conserved regions of the alignment. We then used the resulting trimmed alignment (477 sequences, alignment length: 755) to construct 50 maximum likelihood phylogenies with 1000 bootstraps in IQtree v. 1.6.12 (parameters: –bb 1000; [33]), rooted in the Orco clade. We visualized and annotated the tree with the best likelihood score with iToL v 6.9 [34]. We labeled major OR clades following established subfamily nomenclature [14].

### OR genomic location and structure analyses

To investigate how ORs diversified in both exon structure and location across the genome, we used a custom script to extract genomic coordinates and exon annotations from the GFF file of newly annotated *P. pyralis* ORs using R v. 4.4.1 [35]. We used the R packages ggtree v. 3.12.0 [36], ggplot2 v. 3.5.1 [37], and gggenes v. 0.5.1 [38] to visualize genomic position and exon structure relative to phylogeny. Scripts are available on GitHub: https://selower.github.io/Photinus_pyralis_ORs/.

### RNA extraction and sequencing

To characterize the OR expression, we isolated RNA from the antennae and hind legs of wild male (N= 4) and female (N = 3) *P. pyralis* captured during their active period (within 30 min of sunset) from May 28-30, 2019 in Brasstown, North Carolina (Additional File 2: Table S2). We maintained captive fireflies in individual polystyrene *Drosophila* vials with a piece of moist unbleached coffee filter paper on the bench at ambient light cycle and temperature until preservation in RNA*later* (Ambion; Millipore-Sigma #R0901; Burlington, MA, USA) following manufacturer instructions. One individual was briefly anesthetized with CO_2_ for use in another experiment testing techniques for non-destructive sampling of surface hydrocarbons and returned to its vial for full recovery prior to preservation. We preserved tissues between 1420 and 1710 hours, representing 6.5-4.5 hours prior to onset of flashing activity in the wild, and within one week of capture. Preserved samples were stored at –80 °C until extraction.

We extracted total RNA from dissected antennae and hind legs using TRIzol reagent (Invitrogen; ThermoFisher Scientific #15596026; Carlsbad, CA, USA) following manufacturer instructions, including the addition of glycogen during precipitation to increase RNA yield from the small tissues (GlycoBlue Coprecipitant, Invitrogen; ThermoFisher Scientific #AM9515). Each sample consisted of either both antennae or both hind legs from a single individual. After extraction, we treated samples for potential genomic DNA contamination using an RNA Clean-Up and Concentration Kit (Norgen Biotek Corp. #23600; Thorold, ON, Canada) following manufacturer’s instructions with optional on-column DNase incubation. We assessed sample yield using a Qubit 4.0 Fluorometer with RNA HS Assay (Invitrogen, ThermoFisher Scientific #Q32852). We sent samples with >100 ng of RNA to Novogene (Beijing, China) for quality assessment, followed by low-input Illumina library preparation and sequencing. Due to a previously undescribed hidden break in firefly 28S rRNA that results in a low RIN value (a known, but understudied, phenomenon in arthropods; e.g. [39]), rather than instituting an arbitrary RIN threshold, we assessed RNA quality by manually examining the Bioanalyzer (Agilent 2100; Santa Clara, CA, USA) traces to determine appropriateness for sequencing. All samples chosen for library preparation had a RIN of ≥ 4.8 (mean: 5.2 ± 0.2; Additional File 2: Table S3). Sequencing libraries were prepared at Novogene using the NEBNext Ultra II RNA Library Prep Kit for Illumina (NEB, USA) following manufacturer recommendations. Samples were sequenced to at least 30 million PE 150 bp reads on an Illumina NovaSeq 6000. The raw reads are available on NCBI SRA (BioProject: PRJNA1196372).

### Differential Expression Analysis

To obtain a transcript set that also included a complete set of ORs for differential expression analysis, we downloaded the most recent genome assembly (v1.3; GCF_008802855; [18]) and annotated transcripts from NCBI. We then identified the NCBI-annotated genomic locations that represented the manually-annotated ORs with reciprocal BLAST (parameters: blastn, e-value: 1e-5). Where there was not a one-to-one match between NCBI and manual annotations, we conducted further curation consisting of (i) splitting chimeric transcripts, where the complete open reading frame of another gene was included on the same transcript with an OR, and (ii) adding novel ORs – manually-annotated OR gene models that had no match in the NCBI annotations (Additional File 2: Table S1).

To perform a differential expression analysis, we initially assessed raw RNAseq reads for quality using FastQC v. 0.11.9 [40], with output files concatenated for easier viewing using MultiQC v. 1.14 [41]. We trimmed low-quality reads using Trimmomatic v. 0.39 (default parameters; [42] and then identified and removed potential contaminant reads using Kraken2 v. 2.1.1 [43], using the PlusPF database that includes archaea, bacteria, viral, plasmid, human, protozoa, and fungal taxa (v. 5/17/2021).

We used the cleaned reads for quasi-mapping and quantification to the curated *P. pyralis* GCF_008802855 mRNAs that included the additional manually-annotated ORs using Salmon v. 1.9.0 [44] with the genome as a decoy (parameters: –-validateMappings –-seqBias –-gcBias). We then imported the quantification into R v. 4.2.2 [35] and screened out genes with low expression by filtering the dataset to retain only genes that had a counts per million (CPM) value greater than one in at least three samples (i.e. the size of the smallest experimental group of interest in this study). Subsequently, we transformed the digital counts into continuous values using variance stabilizing transformation (VST) from the DESeq2 R package v. 1.36.0 [45]. We then verified that the major sources of variation in the data could be attributed to our experimental design rather than confounding factors using principal component analysis.

To test for differential expression, we followed the general flow of the lmerSeq methodology and used linear mixed models with random effects to model the VST expression levels [46]. For fixed effects, we included binary sex, binary tissue type (leg vs. antennae), and the interaction between the two, while using a random intercept for each specimen to account for the paired nature of the data (hind legs and antennae were sampled from each individual). Additionally, we utilized the empirical Bayes procedure implemented in the REBEL R v. 0.1.0 [47] package to stabilize the random intercept variance estimates, thus minimizing the number of genes with singular fits that would have to be discarded due to unreliable p-values and regression coefficients. We then used linear contrasts of the regression coefficients to conduct the comparisons of interest, where t-statistics and their associated multiple-comparison-corrected p-values were extracted for inference. Significance was determined at FDR < 0.05. We assessed model fit by examining residuals versus fitted values for all ORs identified as differentially expressed. We used REBEL estimates of differential expression, approximately on the log_2_ scale, and VST counts for subsequent plotting. The scripts and HTML of our data analysis are available on GitHub: https://selower.github.io/Photinus_pyralis_ORs/.

## Results

### 101 ORs + Orco identified

Using an exhaustive iterative blast search strategy with permissive e-value followed by manual curation, we found 101 ORs and the conserved Orco in the published *Photinus pyralis* genome assembly [18], Figure 1). Of these, one OR (PpyrOR8) had two potential splice forms, with mutually exclusive copies of the initial two exons. Two ORs were partial, missing the N-terminus (PpyrOR101NTE) or C-terminus (PpyrOR83CTE), while 15 had premature stop codons and were classified as pseudogenes (PSE, Additional File 2: Table S1). As expected, the majority of annotated ORs were predicted to have seven transmembrane domains (6.73 ± 0.76; TOPCONS; Additional File 1: Text S1, Additional File 2: Table S1). The number of ORs, rarity of alternative splice forms, presence of pseudogenes, and seven transmembrane domain structures are consistent with findings in other beetles [14].

**Figure 1.**
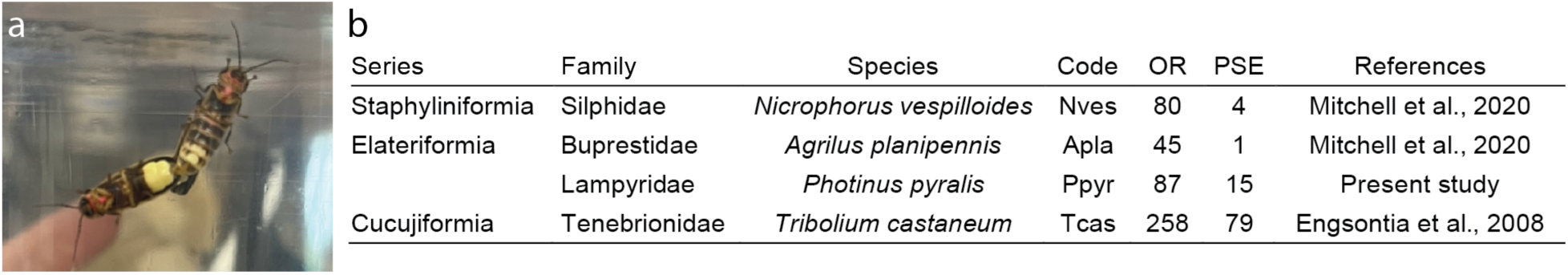
*Photinus pyralis* odorant receptor numbers are in the range of beetles. a) A common eastern firefly (*P. pyralis*) male (left) and female (right) mating in the lab. Photo: Lauren Shaffer. b) Numbers of *P. pyralis* odorant receptors (OR) and OR pseudogenes (PSE) compared to other beetle species used in this study. Code: the four-letter abbreviation used to name ORs in the phylogeny.

### Expansion of Group 2B ORs throughout the genome

A maximum likelihood phylogeny comparing *P. pyralis* ORs with ORs from a subset of beetle species that span OR diversity shows that, in addition to Orco, *P. pyralis* has members of OR Groups 2A, 2B, 3, 4, and 6 (Figure 2). Group 2B is particularly expanded in *P. pyralis*, with all the identified pseudogenes occurring in this clade. Group 2A ORs have the canonical OR structure, with distinct exons A,B,C,D, and E (Figure 3, Additional File 1: Text S2; Additional File 2: Table S4). All other ORs had introns inserted into the A exon, with ORs from Groups 2B, 3, 6, and 4 generally having fragmented A exons, sometimes fused to B exons. The median intron length of full-length ORs was 49 nucleotides (mean: 428 +/-1691, range: 38 – 25,152; Additional File 1: Text S3). None of the *P. pyralis* ORs were closely related to a beetle OR with a known ligand. Interestingly, the phylogeny did not mimic species relationships – *A. planipennis* ORs were generally not sister to *P. pyralis* ORs, except for Orco and OR1. The OR1 clade, consisting of AplaOR1 and PpyrOR1, though nested within the paraphyletic Group 2 clade, is divergent enough to warrant closer examination for classification.

**Figure 2.**
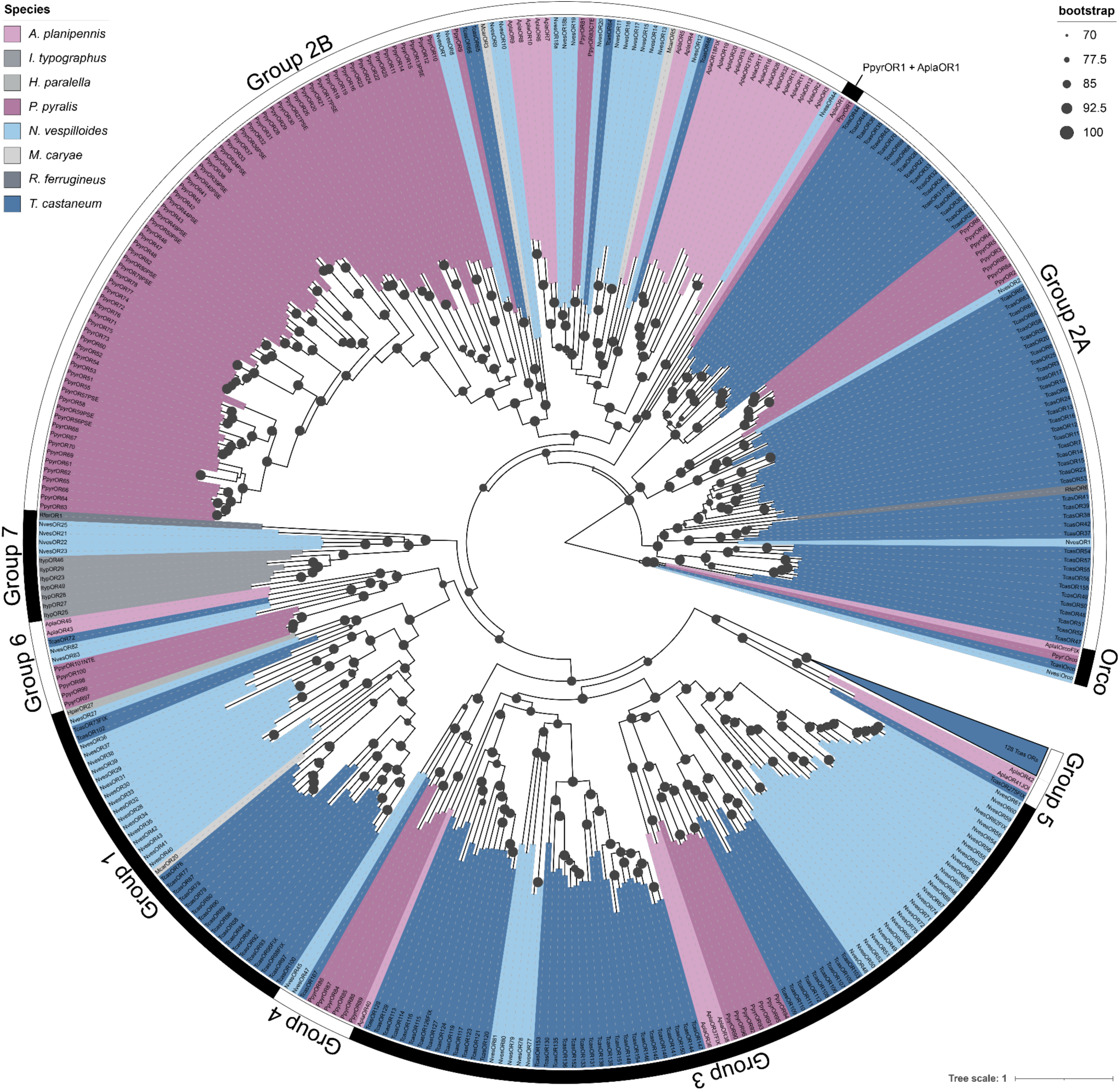
Group 2 ORs expanded in *Photinus pyralis*. A maximum likelihood phylogeny comparing *P. pyralis* ORs (Ppyr; shaded labels) to complete OR repertoires from *Agrilus planipennis* (Apla), *Nicrophorus vespilloides* (Nves), *Tribolium castaneum* (Tcas). Also included are ORs with known ligands from other beetle species: *Megacyllene caryae* (Mcar), *Ips typographus* (Ityp), *Holotrichia parallela* (Hpar), and *Rhynchophorus ferrugineus* (Rfer). Size of circles at nodes corresponds to bootstrap values and clades with less than 70% bootstrap support are collapsed into polytomies. One clade of Group 5A Tcas ORs is depicted as a triangle for visual clarity. OR subfamily (group) identity follows Mitchell et al. 2020. *P. pyralis* has members of Groups 2, 3, 4, and 6. PpyrOR1 is in a divergent clade with AplaOR1 within the paraphyletic Group 2 ORs, suggesting that OR1s require further consideration for classification. PpyrORs are divergent from beetle ORs with known function.

**Figure 3.**
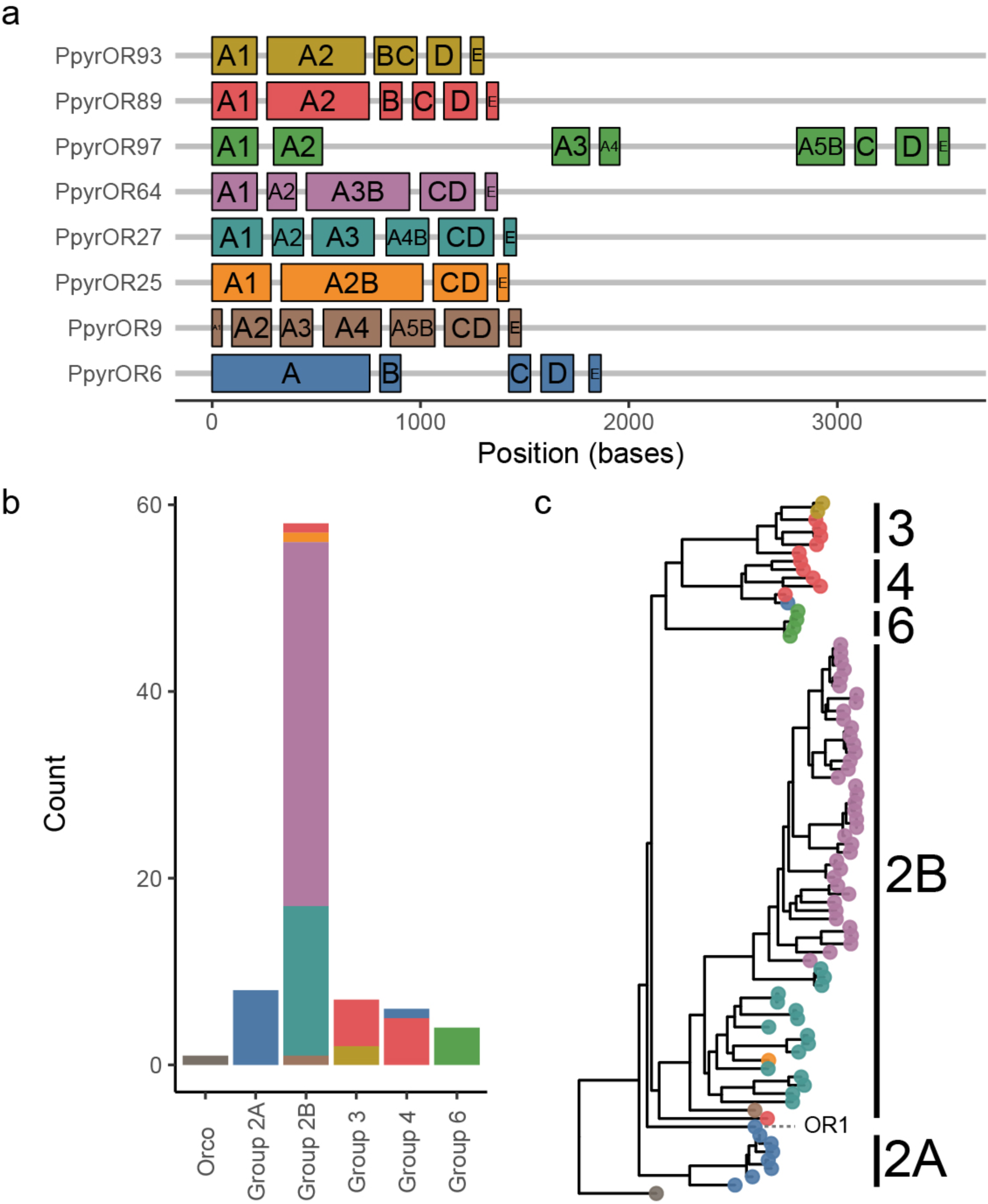
Variation in OR exon structure in *Photinus pyralis*. a) Example exon/intron structures of full-length PpyrORs. We selected the shortest (starting nucleotide to ending nucleotide) OR from each structure type for display (see Additional File 1: Supplementary Text S2 for the full display). All structures are variants of the canonical five-exon (A, B, C, D, E) structure in beetles (14). b) The number of ORs in each OR group, colored by structure as in panel (a). Group 2B has the most structural diversity. c) Exon structures on the PpyrOR phylogeny. Colored circles at tip correspond to structures in (a). Phylogeny from Figure 2, trimmed to only include full-length PpyrORs.

ORs are found on every linkage group (LG) in the *P. pyralis* genome assembly. However, not every OR group is present on every LG. Group 2B has the greatest distribution throughout the genome, as it is found on every LG except for LG10 (Figure 4), with the most occurrences on LG4. This expansion includes one Group 2B OR on LG3a, the putative *P. pyralis* X chromosome. In contrast, Groups 2A and 3 are the most restricted, each found on only a single LG (LG1 and LG10, respectively). All other groups, except for Orco on LG10, are located on at least 2 LGs. While LG4 has the highest absolute number of ORs, most of which are from OR group 2B, LG3b hosts the highest OR group diversity, with members of Groups 2A, 2B, 4, and 6.

**Figure 4.**
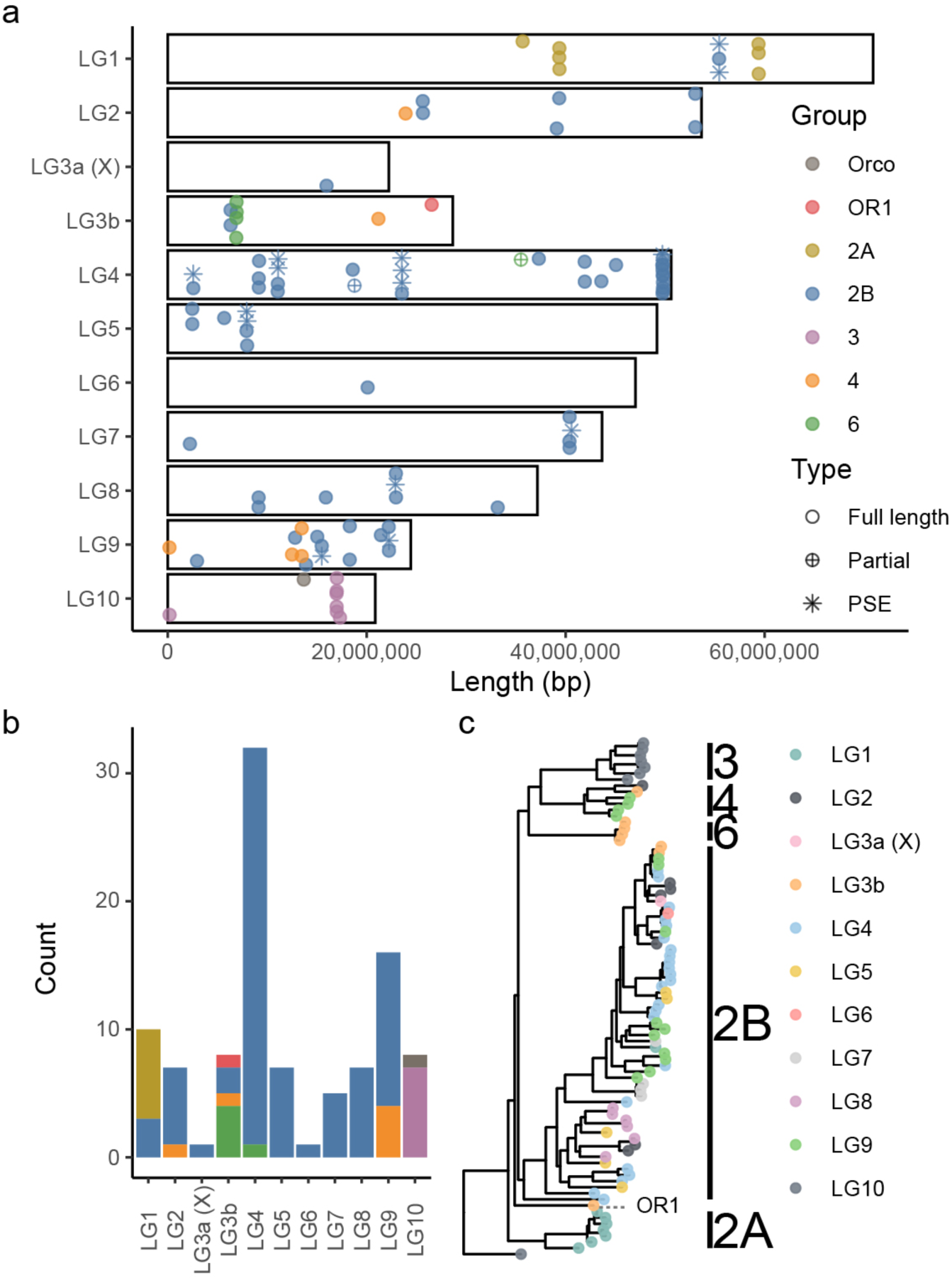
PpyrORs diversified across linkage groups. a) Position of full length, partial, and pseudogenized (PSE) PpyrORs on linkage groups (LG, white bars), colored by OR group. Points are vertically jittered for visualization. b) Number of ORs on each LG, colored by OR group as in (a). LG4 has the highest number of ORs, dominated by 2B ORs. LG3b hosts the most OR diversity (four OR groups). Pseudogenized ORs are on six out of 10 LGs. Group 3 and 2A ORs are limited to a single linkage group, while Group 2B ORs are on every linkage group except LG10, including the sex chromosome (LG3a). c) ORs have diversified across linkage groups (LGs). Colored circles at tips correspond to LGs. Phylogeny is identical to that in Figure 2, trimmed to only include PpyrORs. Some OR groups diversified on the same linkage group (e.g. Group 3 ORs), while others are distributed throughout the genome (e.g. Group 2B).

### A subset of ORs are upregulated in adult antennae

Pseudoalignment of the PE 150 bp Illumina RNA sequence reads from antennae and hind legs of wild-caught signaling *P. pyralis* (4 male, 3 female; 41.1 +/-6.6 million reads per sample after cleaning, Additional File 2: Tables S2, S3) yielded 13,340 genes with a CPM > 1 in at least 3 samples, including 45 of the 101 manually-annotated PpyrORs (Orco, 41 full length ORs, and 3 OR pseudogenes). The remaining ORs may be expressed in other tissues, at different times, or in different life stages.

Principal components analysis (PCA) of the variance stabilized counts revealed that tissue (PC1) and sex (PC2) accounted for 57.1% of the variance in gene expression (45.6% and 11.5%, respectively; Additional File 1: Text S4; Additional File 2: Table S5). Thus, our experimental variables accounted for the major axes of variation in gene expression in our dataset. Unknown factors at capture such as infection status, age, and mating status may have contributed to the remaining PCs.

All 45 ORs that passed filtering had non-zero expression in all tissues of interest. 32 and 36 ORs were significantly upregulated in male and female antennae, respectively, as compared to legs (Figures 5 and 6, Additional File 1: Text S5, S6, Additional File 2: Table S6). 30 of these were upregulated in the antennae of both sexes relative to hind legs, while two were significantly upregulated only in male antennae relative to male hind legs (PpyrOR67, OR100), and six were significantly upregulated only in female antennae relative to female hind legs (PpyrOR15, OR23, PpyrOR28, OR59PSE, OR64, OR81). There were 10 differentially expressed genes between male and female hind legs, none of them ORs.

**Figure 5.**
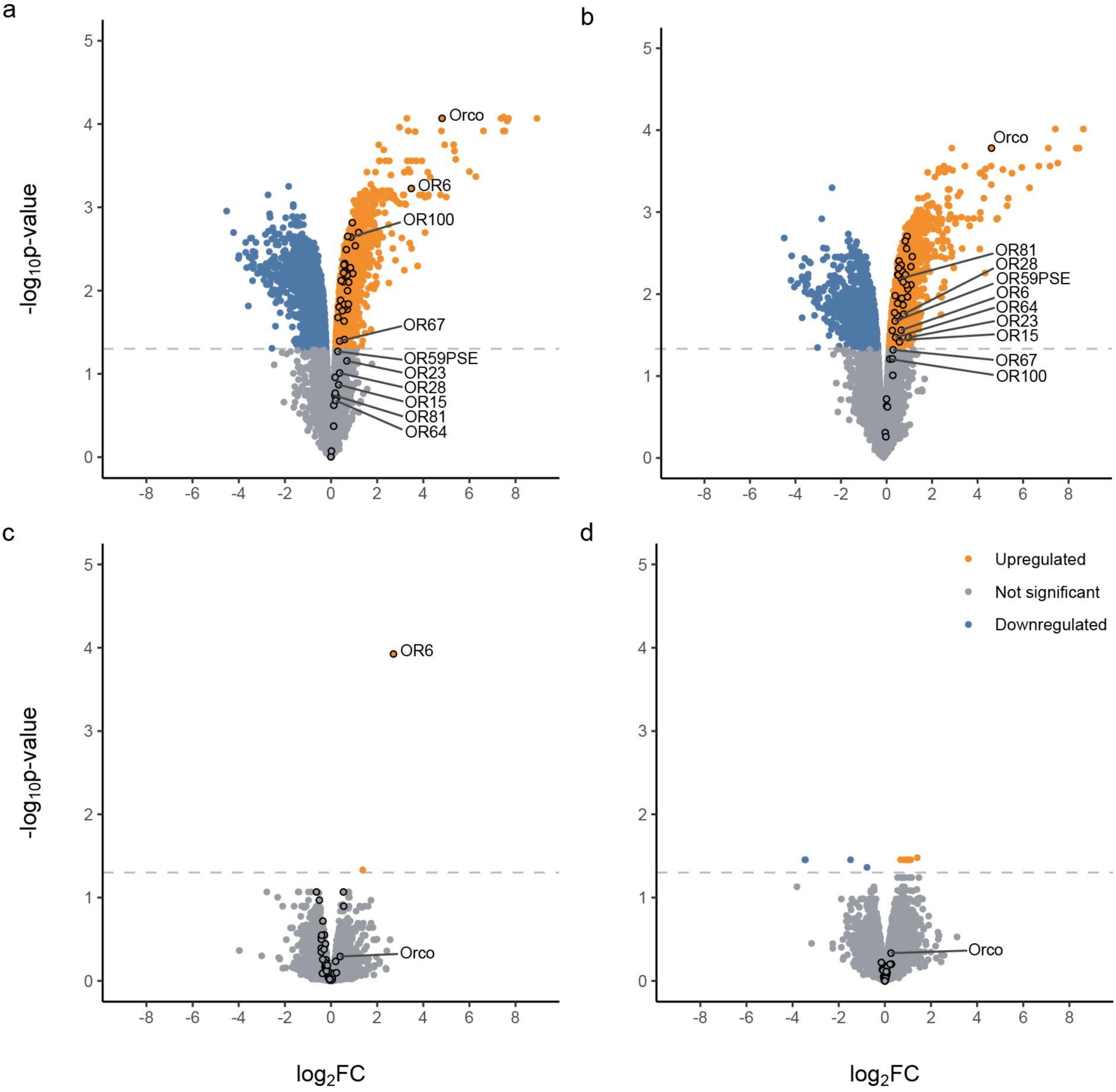
OR expression is antenna-biased. Volcano plots of fold-change versus significance in (a) male antennae versus hind legs, (b) female antennae versus hind legs, (c) male versus female antennae, (d) male versus female hind legs. Inset: Orange points show genes that are significantly upregulated, while blue points show genes significantly downregulated, while grey shows genes that did not reach significance. Black outlines indicate ORs. 32 and 36 ORs are upregulated in males and females, respectively. Orco is significantly upregulated in the antennae of both sexes. PpyrOR6 is the next most highly expressed OR in male antennae (a) and the only differentially expressed gene between male and female antennae (c). A subset of ORs are significantly upregulated in the antennae of only one sex – PpyrOR67 and 100 are significantly upregulated in males, but not in females (a, b) and PpyrOR15, 23, 28, 59PSE, 64, and 81 are significantly upregulated in females, but not in males (a, b).

**Figure 6.**
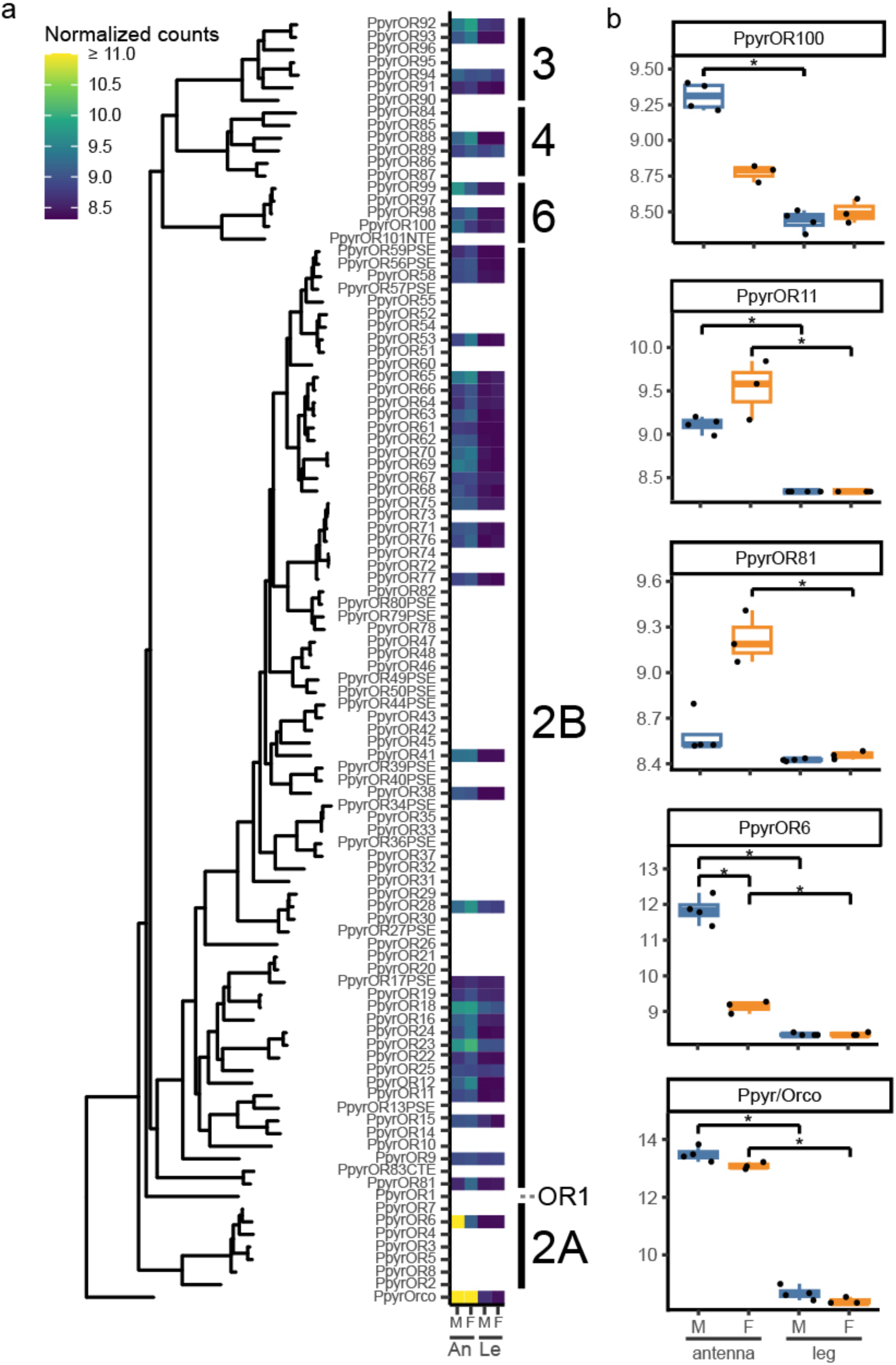
OR expression patterns across the phylogeny. a) Normalized (VST) counts for each OR on the phylogeny presented in Figure 2, trimmed to only *Photinus pyralis* ORs. Bars at right show the OR group designation. M: Male, F: Female, An: Antennae, Le: Hind legs. ORs with blank counts did not pass initial filtering (CPM > 1 in at least 3 samples). b) Boxplots of normalized (VST) counts for each tissue by sex combination for six exemplar ORs, listed in order of placement in the phylogenetic tree (top to bottom): PpyrOR100 (significantly upregulated in antennae vs legs in males only), PpyrOR11 (significantly upregulated in antennae relative to legs in both sexes), PpyrOR81 (significantly upregulated in antennae relative to legs in females only), PpyrOR6 (significantly upregulated in male vs female antennae), and Orco (the most highly expressed OR in antennae). Black jittered points show values for each sample. Blue boxplots show males, while orange boxplots show females. Conditions that are significantly different are indicated by bars, where * = p < 0.05. See Additional File 1: Figure S5 for boxplots of all significantly differentially expressed ORs.

### PpyrOR6 shows sex-biased expression

Two genes were significantly upregulated in male as compared to female antennae (Additional File 2: Table S6). One of these was an OR – PpyrOR6 (Estimate: 2.71, p < 0.05; Figure 5, 6). The other gene was uncharacterized. No ORs were significantly differentially expressed between male and female hind legs.

## Discussion

Here we have identified, for the first time, a comprehensive OR repertoire for a firefly species. Like comparative studies with other beetle ORs, we found lineage-specific diversification in *P. pyralis*, with evidence of gene duplication both in tandem arrays and across the genome. Further, we have shown that ∼38% of ORs are upregulated in adult antennae as compared to hind legs, including a single OR that is significantly upregulated in male antennae.

### Manual annotation yields a comprehensive OR repertoire

The total number of ORs in *P. pyralis* is within the expected range for beetles [14]. Manual curation was effective – we annotated 75 more gene models than using a conservative ortholog search strategy (e-value: 10^-5^, N = 26 ORs + Orco; [48]), and 65 more than NCBI’s automated annotation (N = 36). By annotating ORs in the genome assembly and subsequently using RNAseq to measure expression, we discovered that fewer than 50% of all ORs are detectable in our RNAseq data. This reinforces previous studies demonstrating that OR repertoire derived from RNA sequencing are significant underestimates of total OR repertoire in beetles (e.g. [21,22]). We recommend that future studies of OR diversity should seek to incorporate both genome and transcriptome data for OR discovery.

### ORs diversify in place and around the genome

Previous studies in beetles have noted a pattern of lineage-specific OR diversification with the addition of data from a representative species from each new major group [14].

Similarly, our data showed that there has been clade-specific OR diversification in fireflies, particularly in Group 2B. PpyrOR1 is in a divergent clade with AplaOR1 within the paraphyletic Group 2 ORs, suggesting that OR1s require further consideration for classification.

Three OR groups were not present, and we presume them lost in *P. pyralis* (Groups 1, 5, and 7). Absence of Group 5A ORs was expected, since they appear to have evolved as late as the Bostrichidae [22], and elateriforms, which include fireflies, diverged before that split [24]. However, Group 5B ORs were absent in *P. pyralis*, which is surprising given their presence in the buprestid beetles (e.g. *A. plannipennis*). This may represent a novel loss in the *P. pyralis* lineage. In contrast, ORs in Groups 1 and 7 appear to be absent in *A. planipennis* as well as *P. pyralis*, which are also elateriforms – evidence that loss of these OR groups happened early in elateriform evolution.

Some ORs occurred in tandem arrays on a single linkage group, such as the Group 3 ORs on LG10, suggesting processes like unequal crossing over playing a role in their duplication. In contrast, Group 2B ORs have proliferated throughout the genome, to every linkage group but LG10, suggesting that long-range duplication events, such as TE-mediated transposition, have occurred. Future work will examine repetitive DNA content with respect to OR genomic location.

### Implications of expression pattern for OR function

A previous study found no behavioral evidence for light-using fireflies employing close-range chemosensation during mating [8]. In our study one of only two genes that were differentially upregulated in male as compared to female antennae was an OR, PpyrOR6.

Because *P. pyralis* males land near females and scramble to approach during mating [17], male antennae are usually the first sensory organ directed in contact with a female. Males tap their antennae over the dorsal surface of the female prior to mating, suggesting a potential role of ORs in detecting female-specific compounds, such as cuticular hydrocarbons (CHCs), at close-range. CHCs provide species identity, sex, and mating status information in other insect species [49–52] and several compounds have been described for fireflies [8,53–55]. Close-range chemosensory signals may also function in predator escape, for instance, when *Photuris* firefly females mimic *Photinus* females to lure *Photinus* males for a meal [56]. Though many CHCs are large molecules, they can be detected at distance without any direct contact [57]. PpyrOR6 may therefore be involved in avoiding predation by sensing odorants from these male-specialist predators.

Eight ORs were found upregulated in antennae relative to hind legs in one sex, but not the other. PpyrOR67 and 100 were upregulated in male antennae relative to male legs, but not in female antennae relative to female legs. Despite no significant upregulation between males and females in antennae, it is possible that these ORs are involved in male-specific functions during mating. PpyrORs 15, 23, 28, 59PSE, 64, and 81 were upregulated in female antennae relative to female legs, but not in male antennae relative to male legs. However, like PpyrORs 67 and 100, they were not significantly differentially expressed between female and male antennae. These genes may have female-specific biological functions, such as selecting oviposition sites to avoid harmful microbes [58]. Notably, we annotated PpyrOR59 as a pseudogene. However, during our annotation process, comparison of RNAseq reads with the reference genome suggested the potential for segregating null alleles of other ORs, e.g., PpyrOR23 (Additional File 2: Table S1). It is possible that a functional version of PpyrOR59 will be found by sequencing additional individuals. Alternatively, PpyrOR59 could be recently pseudogenized.

44% of the full OR repertoire did not pass filtering steps in RNAseq analysis, meaning that they were poorly or not expressed in our samples. These ORs are likely expressed at other times or in other life stages. *P. pyralis* spend 1-2 years in their soil-dwelling larval stage where they hunt soft bodied invertebrate prey. Chemosensation likely plays a significant role in larval ecology, which involves navigating low-light environments with a rudimentary larval visual system, including living underground in winter and summer, and nocturnal activity [59,60].

Taken together, our results indicate that *P. pyralis* males may use olfaction for conspecific or heterospecific species recognition during mating. As we sampled only two tissues from adults, future studies could gather evidence for potential function by examining OR expression across additional tissues, times of day, parts of the season, and life stages. Because phylogenetic relationships did not yield clues as to potential ligands, future behavioral and neurophysiological studies will seek to functionally characterize these ORs, particularly PpyrOR6.

### P. pyralis hosts a relatively large OR repertoire

*P. pyralis* maintains a relatively large OR repertoire as compared to buprestids, a related beetle group that also relies primarily on visual signals. This could be because *P. pyralis* uses chemosensation in the adult stage for mate recognition or other activities, or in other life stages. In other beetles, OR number correlates with dietary breadth [21]. While larval *P. pyralis* are considered earthworm specialists with limited evidence for adult *Photinus* feeding [61,62], suggesting some dietary specialization, *P. pyralis* has also been found nectaring on milkweed [63]. It is also possible that this large OR repertoire is a result of phylogenetic inertia, i.e., ORs in *P. pyralis* represent a repertoire similar in size from a common ancestor. This hypothesis is particularly intriguing as *Photinus corruscus*, a diurnal, exclusively pheromone-using firefly, and *P. pyralis* diverged around 25 million years ago [64]. While *P. pyralis* is known primarily for visual bioluminescent mating signals, it is possible that *P. corruscus* was predisposed for evolution of a volatile mating pheromone due to the molecular machinery in place, and perhaps even in use, in their likely lighted common ancestor [5,6]. Both light and pheromone signals are used for mating by the glow-worm firefly *Lamprohiza splendidula* [4] and are also presumed in other species with neotenic females (e.g., *Pyrocoelia rufa*, [2]). Our results suggest that simultaneous and/or sequential use of light and pheromones in firefly courtship is more widespread than previously considered. New firefly genomes (e.g. [65–68]) becoming available will facilitate the future examination of OR repertoire relative to primary mating signal modality across firefly diversity.

## Declarations

### Ethics approval and consent to participate

*Photinus pyralis* is an unregulated invertebrate and is listed as of “Least Concern” by IUCN. All specimens were collected with landowner permission (see Acknowledgements).

### Consent for publication

Not applicable.

### Availability of data and materials

Annotation, alignment, and phylogeny files are available on Figshare: (DOI:10.6084/m9.figshare.28365455). Illumina RNAseq data has been deposited at NCBI SRA (BioProject: PRJNA1196372). Code for analysis and figure generation is available on GitHub: https://selower.github.io/Photinus_pyralis_ORs/.

### Competing interests

The authors declare that they have no competing interests.

### Funding

This study was funded by the University of Wisconsin Oshkosh (to RFM), and Bucknell University (C. Graydon and Mary E. Rogers Fellowship to SEL; Scadden Research Fellowship to SEL, GP, DC), and the National Science Foundation (IOS-2035286 to SEL, DC and IOS-2035239 to GP).

### Author Contributions

SEL conceived, designed, and coordinated the study, acquired data and funding, analyzed and interpreted the data, and took a primary role in drafting and revising the manuscript. SP analyzed and interpreted expression data, drafted relevant parts of the methods, and revised the manuscript. KM analyzed genomic location, interpreted the data, drafted relevant parts of the methods, figure legends, and results, and revised the manuscript. SD conducted transmembrane domain analysis and interpretation, drafted relevant parts of the methods and results, and revised the manuscript. HT collected specimens and extracted RNA, assembled transcriptomes and annotated ORs, drafted the introduction and relevant methods sections, and revised the manuscript. MP annotated ORs and revised the manuscript. BV consulted on RNAseq analysis methods and interpretation, including writing scripts and running data, drafted relevant sections of the methods, and revised the manuscript. GP acquired funding, and revised the manuscript. DC acquired funding, and revised the manuscript. RM annotated ORs, consulted on phylogeny construction and interpretation, and substantially revised the manuscript.

## Acknowledgments

The authors would like to thank Staurolite Farm for permission to collect, Lourenço Martins and Margot Popecki for collection assistance, and Lauren Shaffer for photos. Thank you as well to Anqi Zhang for early work with the *P. pyralis* transcriptome. Many thanks to Jeremy Dreese and Mike Harvey for smooth administration of Bucknell’s High Performance Computing Cluster, BisonNet (NSF #1659397). Heartfelt appreciation to the cohort, coaches, and staff at the Cold Spring Harbor Scientific Writing Retreat 2024 for support and feedback.

## Additional files

### Additional Files

● 1. Supplementary text for ORs in Ppyr
● 2. Supplementary tables for ORs in Ppyr

### *Data files on Figshare* (DOI: 10.6084/m9.figshare.28365455)

● S1.PpyrOR_Nucleotides_FINAL_2025_02_27.fasta
  ○ Nucleotide sequences for PpyrORs in fasta format
● S2.PpyrOR_Nucleotides_FINAL_2025_02_27.gb
  ○ Nucleotide sequences for PpyrORs in genbank format
● S3.PpyrOR_Protein_FINAL_2025_02_27.fasta
  ○ Amino acid sequences for PpyrORs in fasta format
● S4.PpyrOR_Protein_FINAL_2025_02_27.gb
  ○ Amino acid sequences for PpyrORs in genbank format
● S5.PpyrOR_Scaffolds_FINAL_2025_02_27.gff
  ○ The gff file of scaffolds used for assessing location and intron/exon structure
● S6.Queries_Agla-Apla-Hred-Nves-Pser-RdomORs.fasta
  ○ Beetle ORs used as queries to identify putative PpyrORs
● S7.PpyrORs_Nves_Apla_Tc_functional_beetles_over100aa_n477_noPSE_noPA R_MUSCLE_no_ambigs_noisyclean_reordered_2025_03_04.fasta
  ○ Alignment with Ppyr and other beetle ORs that generated the phylogeny in Figure 2
● S8.run_2.contree_rooted_2025_03_11.newick
  ○ Consensus maximum likelihood phylogeny from IQtree, rooted in Orco
● S9.renumbered_Ppyr_tx_2025_03_04.fna
  ○ Transcripts used for salmon analysis, edited to incorporate the gene models described in this study

## Notes

### Competing Interest Statement

The authors have declared no competing interest.

