## Additional File 1: Supplementary Text for "Antennal RNAseq reveals odorant receptors with sex-biased expression in the common eastern firefly, *Photinus pyralis*"

Supplementary text for ORs in *P. pyralis*

S1. A majority of *Photinus pyralis* odorant receptors have seven transmembrane domains

Distribution of transmembrane domain (TMD) counts of *Photinus pyralis* ORs, as predicted by TOPCONS, a membrane topology prediction webserver (Tsirigos et al. 2015). ORs are categorized as either full length (green), CTE (missing one or more exons on the C-terminus; red), NTE (missing one or more exons on the N-terminus; blue), or PSE (pseudogenes; purple).

The majority of *P. pyralis* ORs have seven transmembrane domains, with an overall range of three to seven TMDs. Full length ORs possess five to seven TMDs, with all NC and NTE ORs possessing four TMDs and pseudogenes possessing either three, six, or seven TMDs. Because insect odorant receptors have seven transmembrane domains (Benton et al. 2006; Smart et al. 2008), the finding that most full length *P. pyralis* ORs are predicted to have six or seven TMDs supports that these gene models are fully functional. In contrast, CTE and NTE PpyrORs were predicted to have only four TMDs, suggesting that CTE and NTEs are not fully functional.

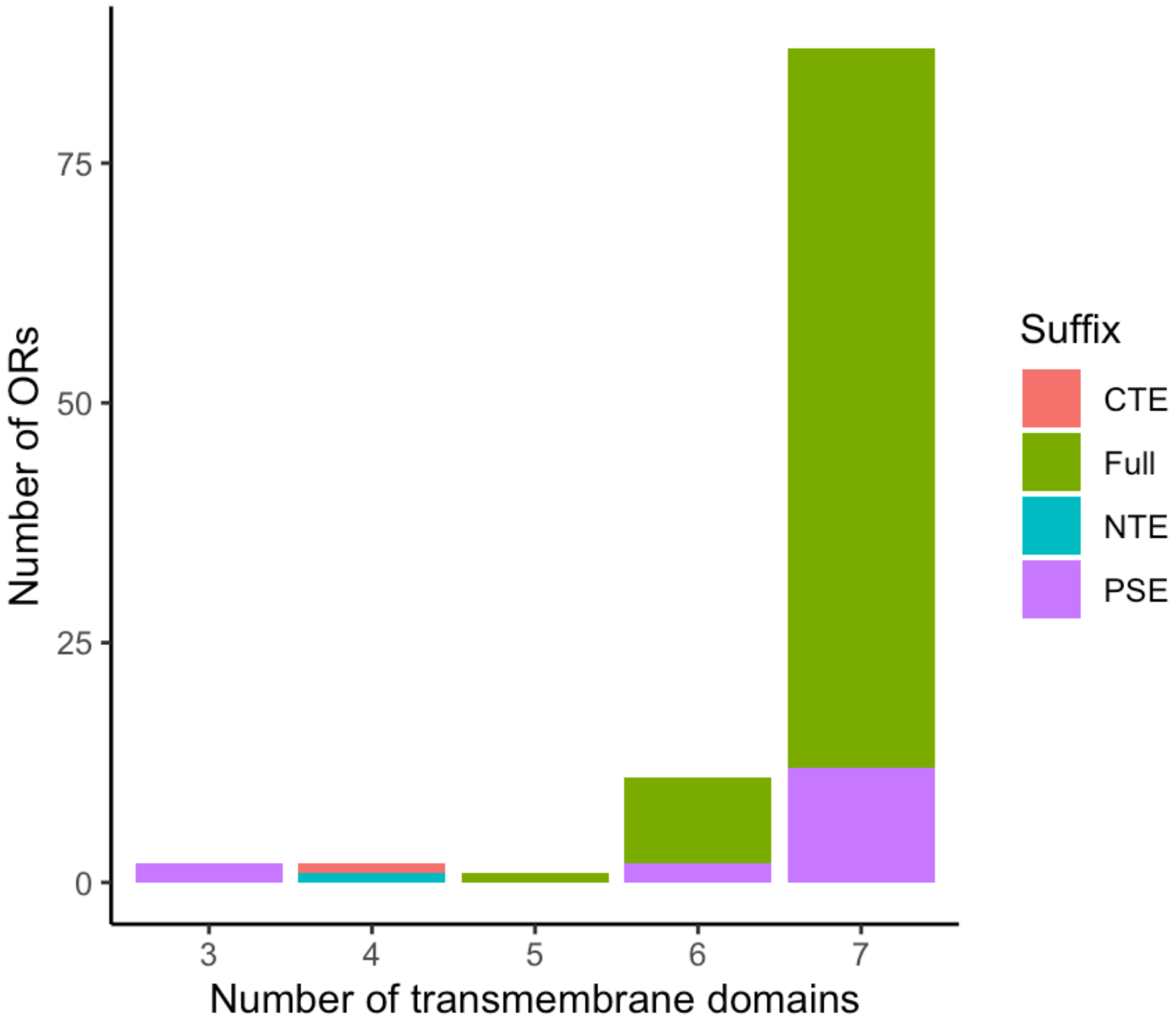

S2. Exon structure across the PpyrOR phylogeny

Phylogeny from main text Figure 2, trimmed to only include full-length PpyrORs. Visualized in R v. 4.4.1 (R Core Team 2020) using the ggtree v. 3.12.0 (Yu et al. 2017) and gggenes 0.5.1 (Wilkins and Kurtz 2023) packages.

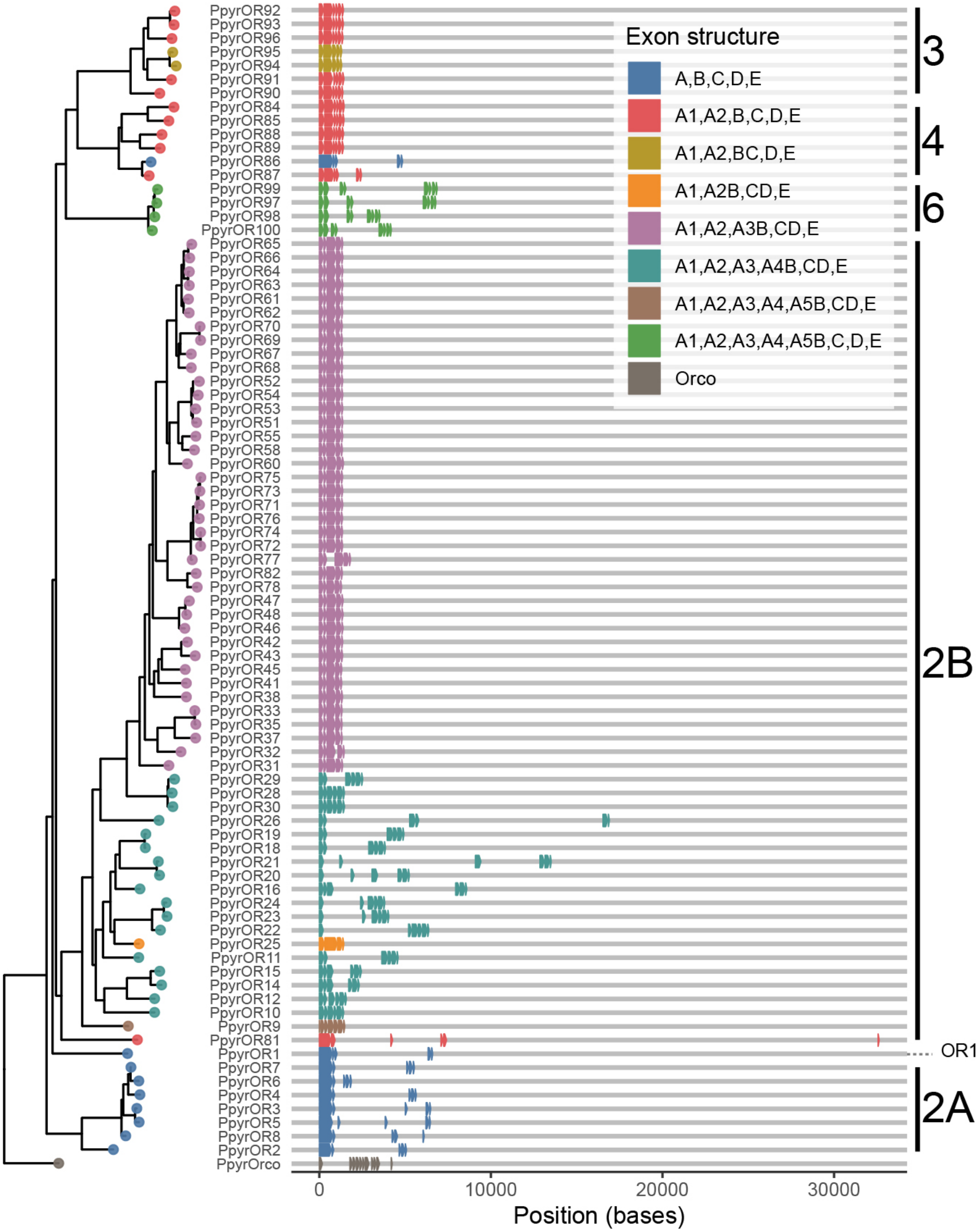

S3. Most PpyrOR introns are short

Histogram of intron length (in bases) across full-length PpyrORs. The median is 49 bases (range: 38 - 25,152 bases).

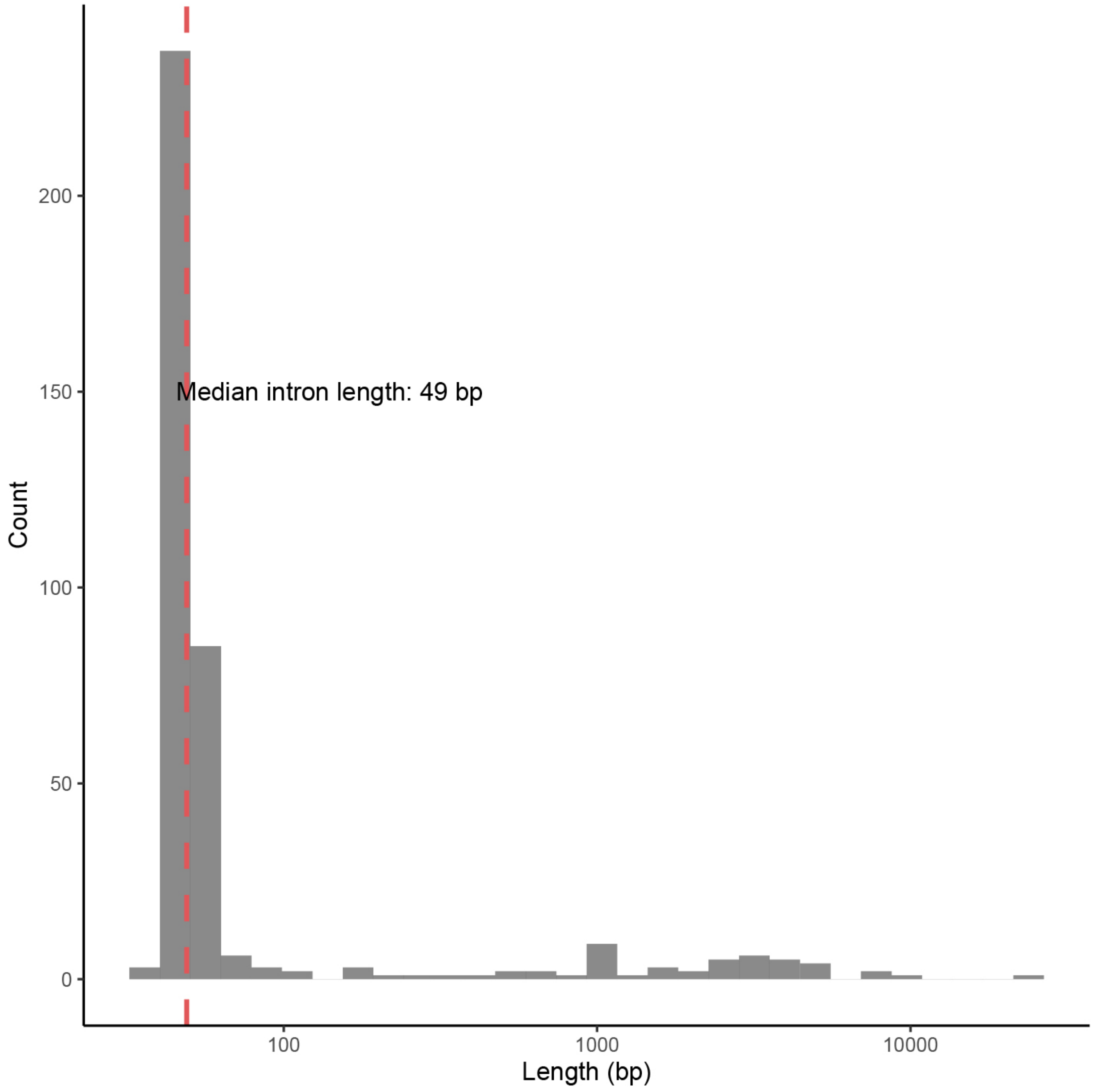

S4. Tissue and sex account for >50% of the variance in gene expression

Principal component analysis (PCA) plot shows the percent variance explained by the top two principal components for the gene expression differences of the samples. Tissue (PC1) and sex (PC2) account for over 50% of the variance in the gene expression of the samples. Fill indicates sample tissue type (gray for antenna and white for leg) and shape denotes sample sex (○ for female and △ for male).

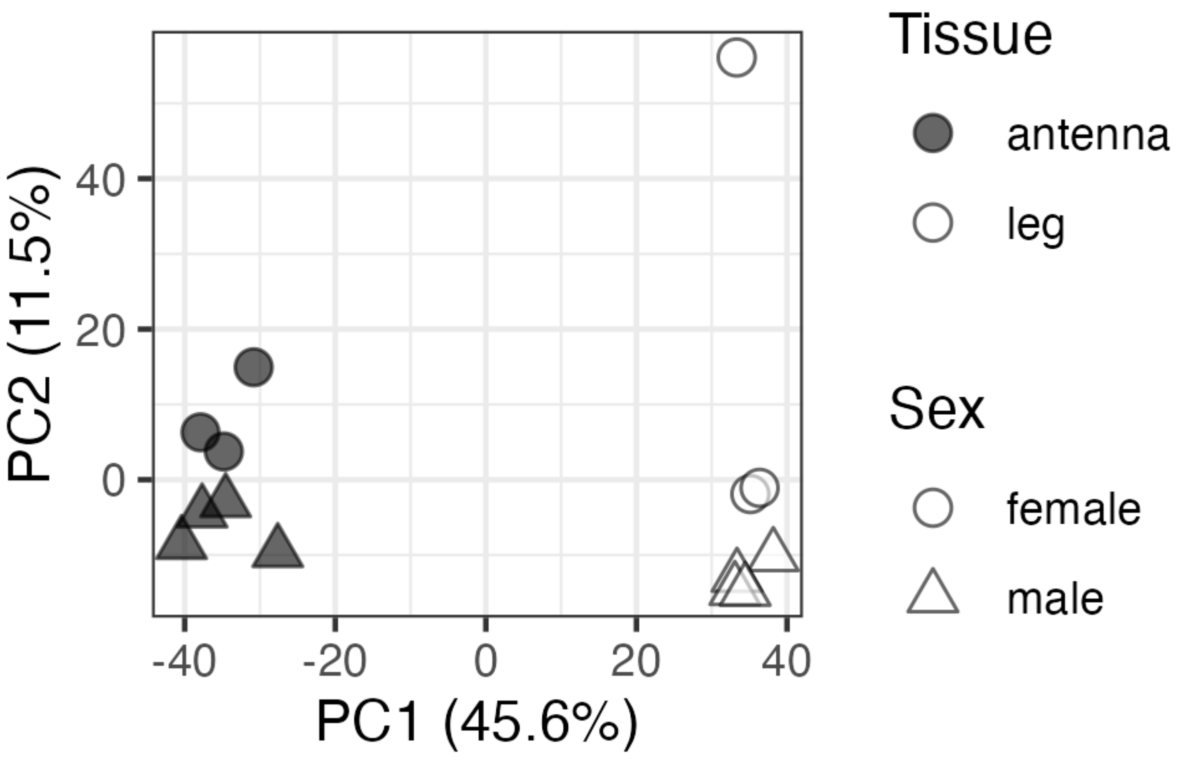

S5. OR expression boxplots

Boxplots of normalized (VST) counts for each tissue by sex combination. Black jittered points show values for each sample. Blue boxplots show males, while orange boxplots show females. Conditions that are significantly different are indicated by bars, where * = $p < 0.05$. Shading of the plot heading shows genes that are significantly differentially expressed in male antennae vs hind legs (dark gray), female antennae v. hind legs (light gray), or both (medium gray); a bold heading outline indicates significantly differential expression between male and female antennae.

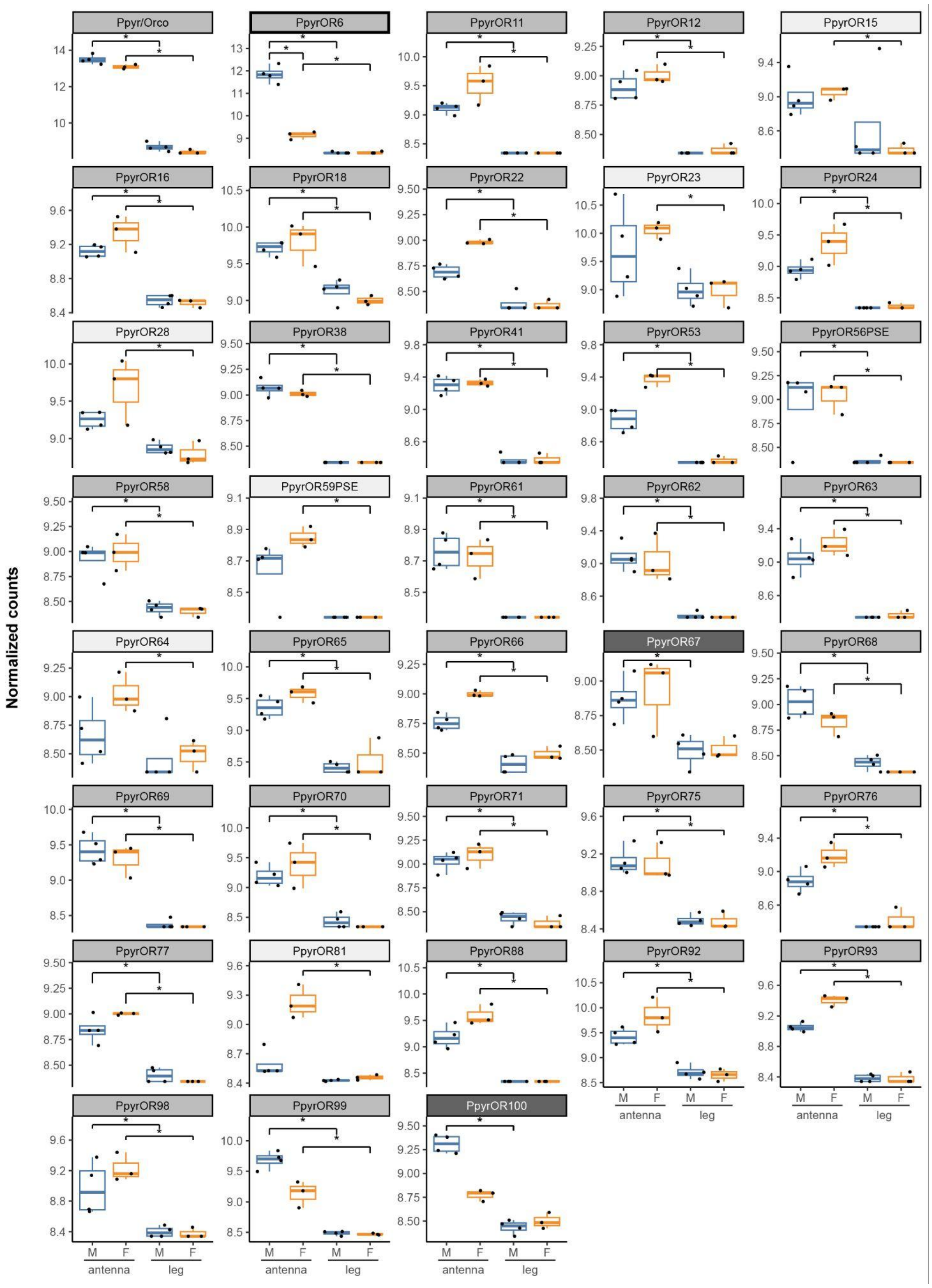

S6. OR expression on linkage groups

Normalized (VST) counts of each OR (circles) on linkage groups (LGs) for a) male antennae, b) female antennae, c) male hind legs, d) female hind legs. Small empty circles show ORs that did not pass initial filtering (CPM > 1 in at least 3 samples). Orco and PpyrOR6, which have the highest normalized counts, are labeled in panel a.

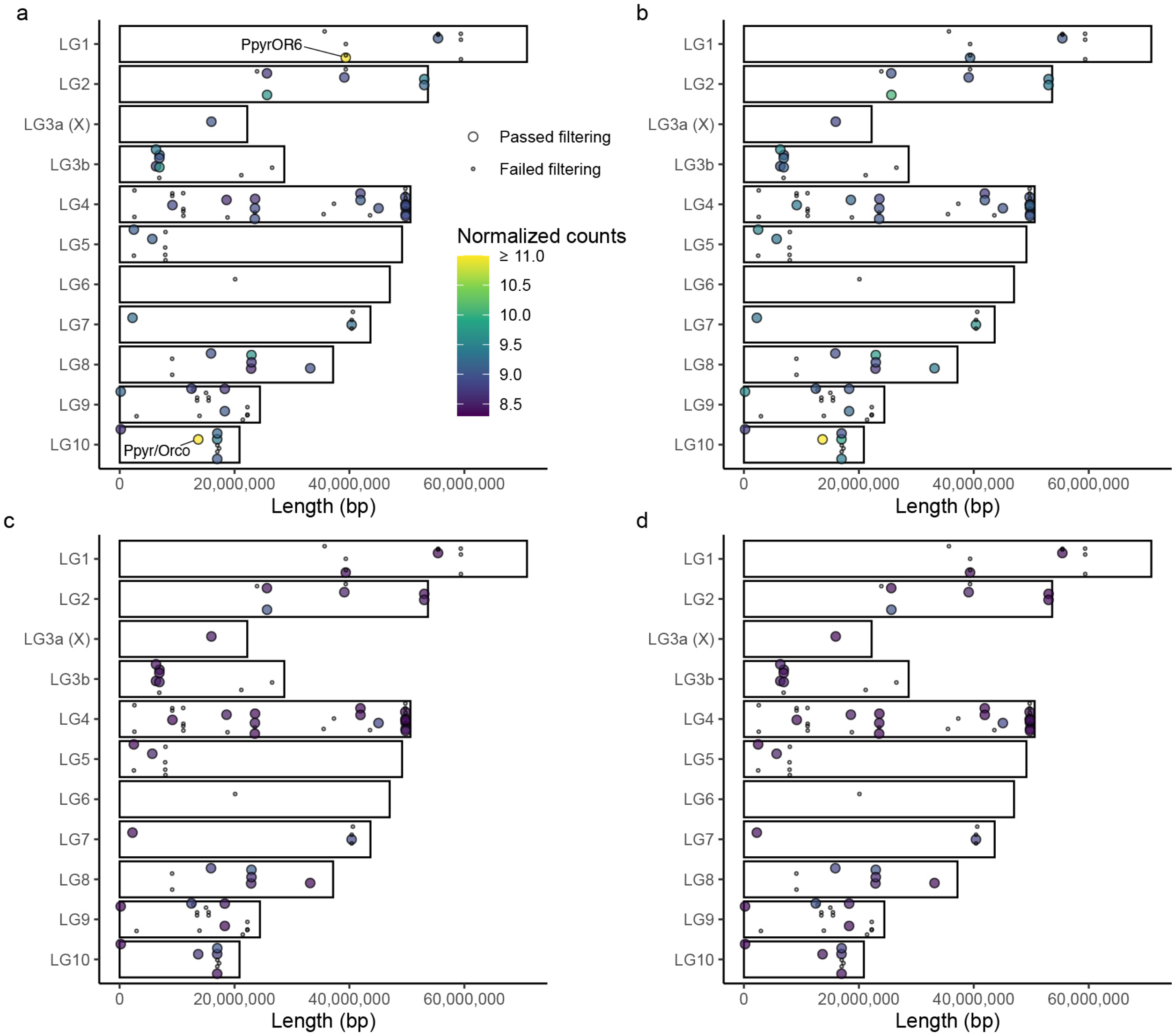
